# Golgi SM protein Sly1 promotes productive *trans*-SNARE complex assembly through multiple mechanisms

**DOI:** 10.1101/2020.01.16.909630

**Authors:** M. Duan, G. Gao, D.K. Banfield, A.J. Merz

**Author notes:** Corresponding author ·.

## Abstract

SNARE chaperones of the Sec1/mammalian Unc-18 (SM) family have critical roles in SNARE-mediated membrane fusion. Using SNARE and Sly1 mutants, and a new *in vitro* assay of fusion, we separate and assess proposed mechanisms through which Sly1 augments fusion: (*i*) opening the closed conformation of the Qa-SNARE Sed5; (*ii*) close-range tethering of vesicles to target organelles, mediated by the Sly1-specific regulatory loop; and (*iii*) preferential nucleation of productive *trans*-SNARE complexes. We show that all three mechanisms are important and operate in parallel, and we present evidence that close-range tethering is particularly important for *trans*-complex assembly when *cis*-SNARE assembly is a competing process. In addition, the autoinhibitory N-terminal Habc domain of Sed5 has at least two positive activities: the Habc domain is needed for correct Sed5 localization, and it directly promotes Sly1-dependent fusion. Remarkably, “split Sed5,” with the Habc domain present only as a soluble fragment, is functional both *in vitro* and *in vivo*.

## INTRODUCTION

SNARE-mediated membrane fusion is central to secretory cargo transport, exocytosis, and organelle biogenesis and homeostasis (Jahn and Fasshauer, 2012; Ungar and Hughson, 2003). Fusion is preceded by tethering, mediated by a diverse group of proteins and usually controlled by small G proteins of the Rab, Arf, or Rho families (Angers and Merz, 2011; Bombardier and Munson, 2015; Pfeffer, 2017; Stenmark, 2012). Tethering is followed by docking: the assembly of a parallel, tetrahelical *trans*-SNARE complex (“SNAREpin”) that links the two membranes (Hanson et al., 1997; Nichols et al., 1997; Sutton et al., 1998; Weber et al., 1998). “Zippering” of the incipient *trans*-SNARE complex does the mechanical work of driving the membranes together to initiate fusion (Zorman et al., 2014). Each *trans*-SNARE complex contains four α-helices, one from each of four SNARE subfamilies: R, Qa, Qb, and Qc (Fasshauer et al., 1998). R-SNAREs often correspond to vesicle- or v-SNAREs, while Qa-SNAREs (also called syntaxins) typically correspond to target membrane, or t-SNAREs. All Qa-SNAREs have in common an N-terminal regulatory “Habc” domain that folds into a trihelical bundle. In some but not all cases the Habc domain can fold back onto the catalytic SNARE domain to form an autoinhibited “closed” conformation (Demircioglu et al., 2014; Dulubova et al., 1999; Dulubova et al., 2001; Fernandez et al., 1998; Kosodo et al., 1998; Misura et al., 2000; Munson and Hughson, 2002; Nicholson et al., 1998; Struthers et al., 2009).

In addition to SNAREs and tethering factors, proteins of the Sec1/mammalian Unc-18 (SM) family have critical roles in SNARE-mediated fusion (Carr and Rizo, 2010; Rizo and Sudhof, 2012; Sudhof and Rothman, 2009). The first SM proteins identified through genetic screens were Vps33a (*carnation* in *Drosophila*) and *Saccharomyces cerevisiae* Sec1 (UNC-18 in *Caenorabditis elegans*; Munc18-1 or nSec1 in mammals; Novick et al., 1979; Patterson, 1932). Despite their early identification and clear importance, and despite major efforts by many laboratories, the general mechanisms of SM function are only now emerging. All SM proteins exhibit strong evolutionary and structural homology, but they interact with cognate SNARE proteins in very different ways. For example, yeast Sly1, yeast Vps45, and Munc18-1 all interact with short N-peptides at the amino termini of their cognate Qa-SNARE proteins (Bracher and Weissenhorn, 2002; Carpp et al., 2006; Dulubova et al., 2002; Furgason et al., 2009; Grabowski and Gallwitz, 1997; Yamaguchi et al., 2002). In contrast, Qa-SNARE N-peptide interactions do not occur with human or yeast Vps33, or with yeast Sec1 (Baker et al., 2015; Dulubova et al., 2001; Lobingier and Merz, 2012; Togneri et al., 2006). We have called SM proteins that interact with Qa-SNARE N-peptides Class I, and those that do not, Class II (Lobingier and Merz, 2012).

Early structural and biochemical studies revealed that Munc18-1 tightly binds the Qa-SNARE Syntaxin-1A in its closed conformation, suggesting an inhibitory role for Munc18-1. (Dulubova et al., 1999; Misura et al., 2000; Yang et al., 2000). However, the emerging consensus is that the core and evolutionarily conserved role of SM proteins is positive, rather than inhibitory. Specifically, SM proteins are hypothesized to nucleate and stabilize fusion-competent *trans*-SNARE complexes (Carr and Rizo, 2010; Sudhof and Rothman, 2009; Toonen and Verhage, 2003; Yoon and Munson, 2018). A breakthrough was achieved in 2015 with two structures of yeast Vps33: one with a Qa-SNARE domain bound, and another with an R-SNARE domain bound (Baker et al., 2015, #55487). When superimposed these structures implied that Vps33 templates the initial assembly of the *trans*-SNARE complex, allowing the *trans*-complex to transit into a metastable “half-zipped” intermediate. Single-molecule force spectroscopy experiments strongly supported this interpretation (Jiao et al., 2018; Ma et al., 2015). It remains unclear whether the templating mechanism is generalizable to all SMs, and it is unclear to what extent SMs promote SNARE-mediated fusion through additional general or pathway-specific mechanisms.

We have turned our attention to Sly1, the ER-Golgi SM. Sly1 has been proposed to promote fusion through several different mechanisms. *First*, meticulous solution biochemistry demonstrated that Sly1 can open the inactive closed conformation of the cognate Qa-SNARE Sed5. This in turn allows Sed5 to more readily complex with Qb, Qc, and R-SNAREs (Bos1, Bet1, and Sec22; (Demircioglu et al., 2014; Kosodo et al., 1998). A limitation of this work is that the SNAREs used were soluble fragments; the roles of Sed5 inhibition and opening were not tested in experiments that assayed membrane fusion. *Second*, we demonstrated that Sly1 binding to quaternary SNARE complexes in solution slows the kinetics of ATP-dependent disassembly by Sec17 and Sec18 (in mammals, α-SNAP and NSF). Consistent with these findings, *in vivo* genetic tests revealed that Sec17 overproduction is tolerated in a wild-type genetic background but becomes lethal when Sly1 function is partially compromised (Lobingier et al., 2014). *Third*, on the basis of structural homology to Vps33, Baker *et al*. (2015) proposed that Sly1 can template Qa- and R-SNARE *trans*-complex formation. *Fourth*, experiments in a companion manuscript (Duan et al., submitted) demonstrate that Sly1 can promote close-range vesicle tethering through an amphipathic helix, α21, that directly interacts with the vesicle membrane.

All of these mechanisms are plausible, yet no study to date has attempted to assess their functional contributions within a unified experimental framework. Here we begin that effort, combining *in vivo* genetic tests with a new chemically defined *in vitro* reconstitution of fusion on the ER-Golgi anterograde pathway. Focusing on Sly1 interactions with the Qa-SNARE Sed5, we demonstrate that multiple mechanisms do indeed contribute to Sly1’s fusion-promoting activity and, unexpectedly, that the regulatory Habc domain of the Qa-SNARE Sed5 augments Sly1-stimulated fusion, rather than being solely auto-inhibitory.

## RESULTS

### Sed5 N-peptide is essential for viability and efficient membrane fusion

The Qa-SNARE Sed5 has four domains: an N-peptide of 21 residues that binds tightly to Sly1; a trihelical Habc domain that is auto-inhibitory; the Qa-SNARE domain; and a C-terminal transmembrane segment (**Fig. 1A**). In a previous study, missense mutations that reduced the affinity of Sly1 for the N-peptides of its client Qa-SNAREs (Sed5 and Ufe1) resulted in minimal defects in assays for viability, secretion, and Sly1 localization. These results were interpreted to indicate the functional “irrelevance” of Sly1-Sed5 interactions (Peng and Gallwitz, 2004). In contrast to these findings, overexpression of a Sly1 cognate N-peptide, or of the Sly1 N-peptide binding domain, shattered the Golgi in mammalian cells (Dulubova et al., 2003; Yamaguchi et al., 2002). To reassess whether the Sed5 N-peptide is functionally important in budding yeast we engineered an allele, *sed5ΔN*, that encodes a Sed5 variant lacking the N-peptide (as defined by the crystal structure PDB 1MQS; Bracher et al., 2002; **Fig. 1A**). In a genetic background expressing wild type *SLY1*, *sed5ΔN* was a recessive lethal allele (**Fig. 1B**; **Supplementary Fig. S1**). Thus, the Sed5 N-peptide is essential for viability. These results are buttressed by recent experiments showing that *sed5*(DFTV), a quadruple missense mutant that impairs Sly1 binding to the Sed5 N-peptide, is also recessive lethal (Gao and Banfield, 2019).

**Fig. 1.**
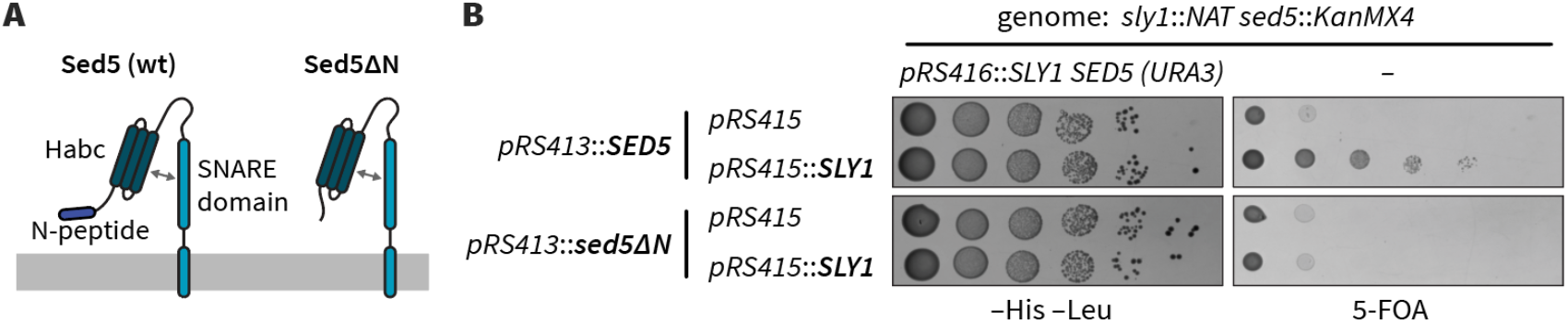
The N-peptide of Sed5 is essential for viability. **A.** Diagram showing the constructs tested. Sed5-ΔN lacks the first 21 aminoacyl residues. **B.** Viability tests. The *sed5-ΔN* allele was tested using *sed5Δ sly1Δ* double knockout cells that carry intact copies of both *SED5* and *SLY1* on a single counter-selectable plasmid. Forced ejection of the *SED5 SLY1* plasmid by plating onto 5-fluoroorotic acid (5-FOA) resulted in lethality unless both *SED5* and *SLY1* were supplied *in trans*. Additional controls are presented in Supplementary Fig. S1.

To test whether the Sed5 N-peptide has a direct role in Sly1-dependent fusion we used a chemically defined assay of fusion driven by ER-Golgi SNAREs (Duan et al., submitted; Zucchi and Zick, 2011). Briefly, we prepare reconstituted proteoliposomes (RPLs) bearing ER-Golgi SNAREs, with two orthogonal FRET reporter pairs. These reporters simultaneously monitor lipid and content mixing in a single 20 μL reaction. Here, we present content mixing results (the reaction endpoint). Fusion requires the presence of Sly1 and also depends on 3% polyethylene glycol, a molecular crowding agent that mimics the action of tethering factors (Duan et al., submitted; Furukawa and Mima, 2014; Lentz, 2007; Mitchison, 2019; Yu et al., 2015). Sly1-mediated fusion is further stimulated by the universal SNARE disassembly chaperones Sec17 (α-SNAP), Sec18 (NSF), and Mg^2+^·ATP (Duan et al., submitted).

To test the function of the N-peptide we prepared RPLs bearing either wild-type Sed5 or Sed5ΔN (lacking the first 21 aminoacyl residues), as well as the Qb- and Qc-SNAREs Bos1 and Bet1. These Q-SNARE RPLs were assayed for their ability to fuse with RPLs bearing the R-SNARE Sec22. In reactions containing Sec17, Sec18, and Mg^2+^·ATP, fusion was rapid and efficient when both 3% PEG and Sly1 were present. However, both the rate and extent of fusion were dramatically reduced when RPLs bearing Sed5ΔN were tested. Moreover, high concentrations of Sly1 were required to stimulate fusion of Sed5ΔN RPLs. With wild type Sed5 near-maximal fusion was observed at 100 nM Sly1, while with Sed5ΔN the rate and extent of fusion were much lower even at 1600 nM Sly1 (compare **Figs. 2C** and **2D**). When PEG(which promotes vesicle tethering) was omitted (**Fig. 2F**), fusion with Sed5ΔN was undetectable even when 1600 nM Sly1 was added. We conclude that the Sed5 N-peptide strongly promotes Sly1-dependent fusion, both *in vivo* and *in vitro*.

**Fig. 2.**
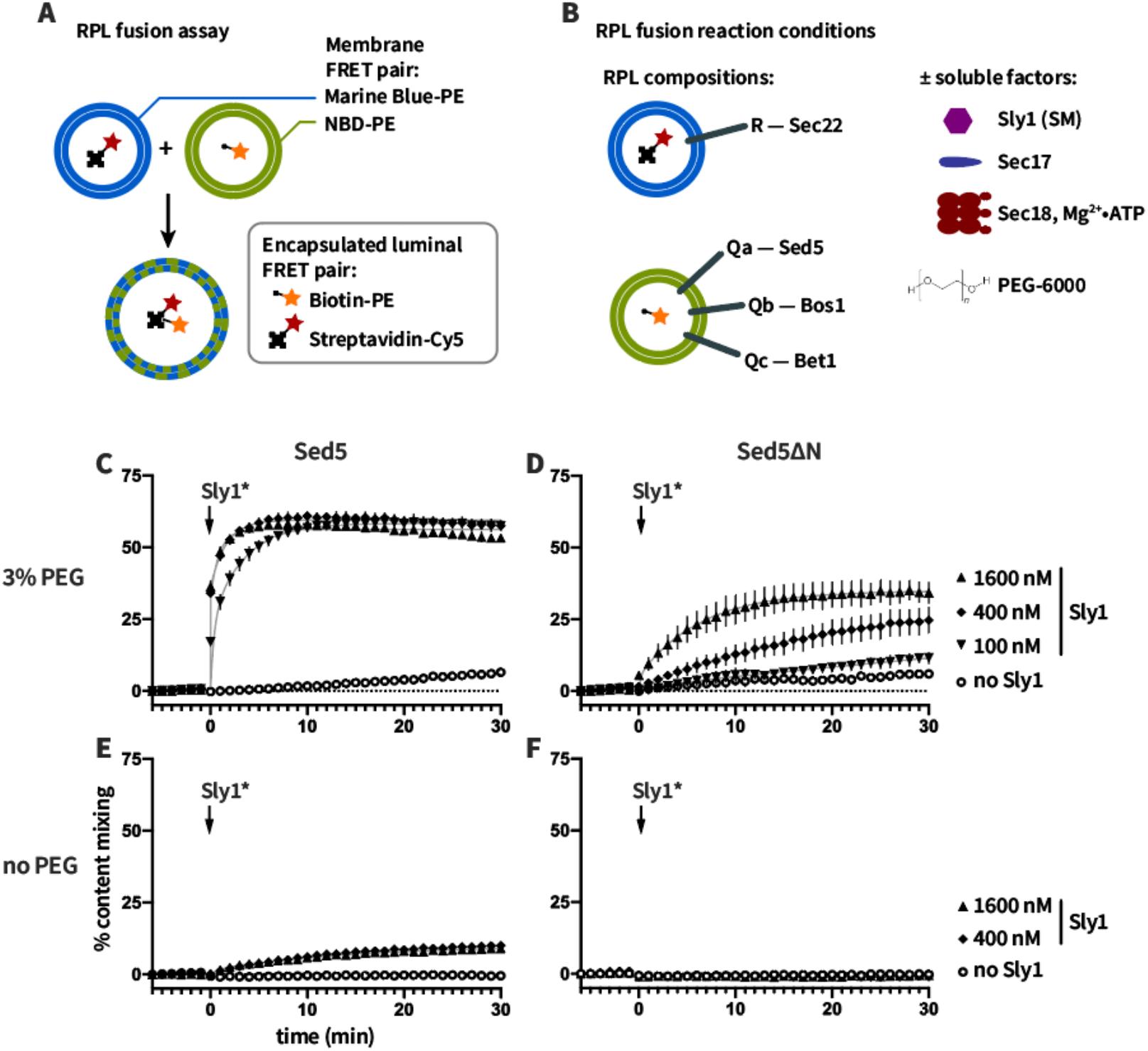
Sed5 N-peptide promotes fusion *in vitro*. **A**. Reporter systems for lipid and content mixing. RPLs are prepared with encapsulated content mixing FRET pair, and with the membranes doped with an orthogonal FRET pair. **B**. SNARE topology and soluble factors added in these experiments. **C-F**. Reactions were set up with RPLs, Sec17, Sec18, Mg2+·ATP, and 3% (C,D) or 0% (E,F) PEG. Q-SNARE liposomes bore either wild-type Sed5 (C,E) or Sed5ΔN (D,F). The reactions were incubated for 5 min and fusion was initiated by adding Sly1 as indicated at t = 0. Points show mean ±sem of at least three independent experiments. Gray lines show least-squares fits of a second-order kinetic function.

### Sly1 hyperactivity suppresses lethality of *sed5ΔN* and restores fusion

The hyperactive allele *SLY1-20* was initially identified as a dominant, single-copy suppressor of loss of Ypt1, the yeast Rab1 homolog (Dascher et al., 1991; Ossig et al., 1991). Single-copy *SLY1-20*, or multi-copy wild type *SLY1*, were subsequently found to suppress deficiencies of a wide variety of proteins that mediate intra-Golgi, ER-Golgi, and Golgi-ER vesicle docking and fusion. In the companion paper (Duan et al., submitted) we show that the mechanism of Sly1-20 hyperactivity involves both release of Sly1 autoinhibition and coupled activation of a latent vesicle tethering activity within the Sly1 auto-inhibitory loop. Somewhat surprisingly, *SLY1-20* was able to suppress the lethal phenotype of *sed5ΔN*, but only when *SLY1-20* was provided on a multiple-copy plasmid (**Fig. 3A**). Single-copy *SLY1-20*, or multiple copy wild-type *SLY1*, were unable to rescue the growth of *sed5ΔN* mutant cells. Thus, the *sed5ΔN* allele is even more deleterious than the already-lethal *ypt1-3* (Rab1-deficient) or *uso1* (p115/Uso1 tethering factor-deficient) alleles — both of which are efficiently suppressed by single-copy *SLY1-20*. The survival of *sed5ΔN* cells containing high-copy *SLY1-20* also allowed us to verify that Sed5ΔN is present at normal abundance and migrates as expected on SDS-PAGE gels (**Fig. 3B**).

**Fig. 3.**
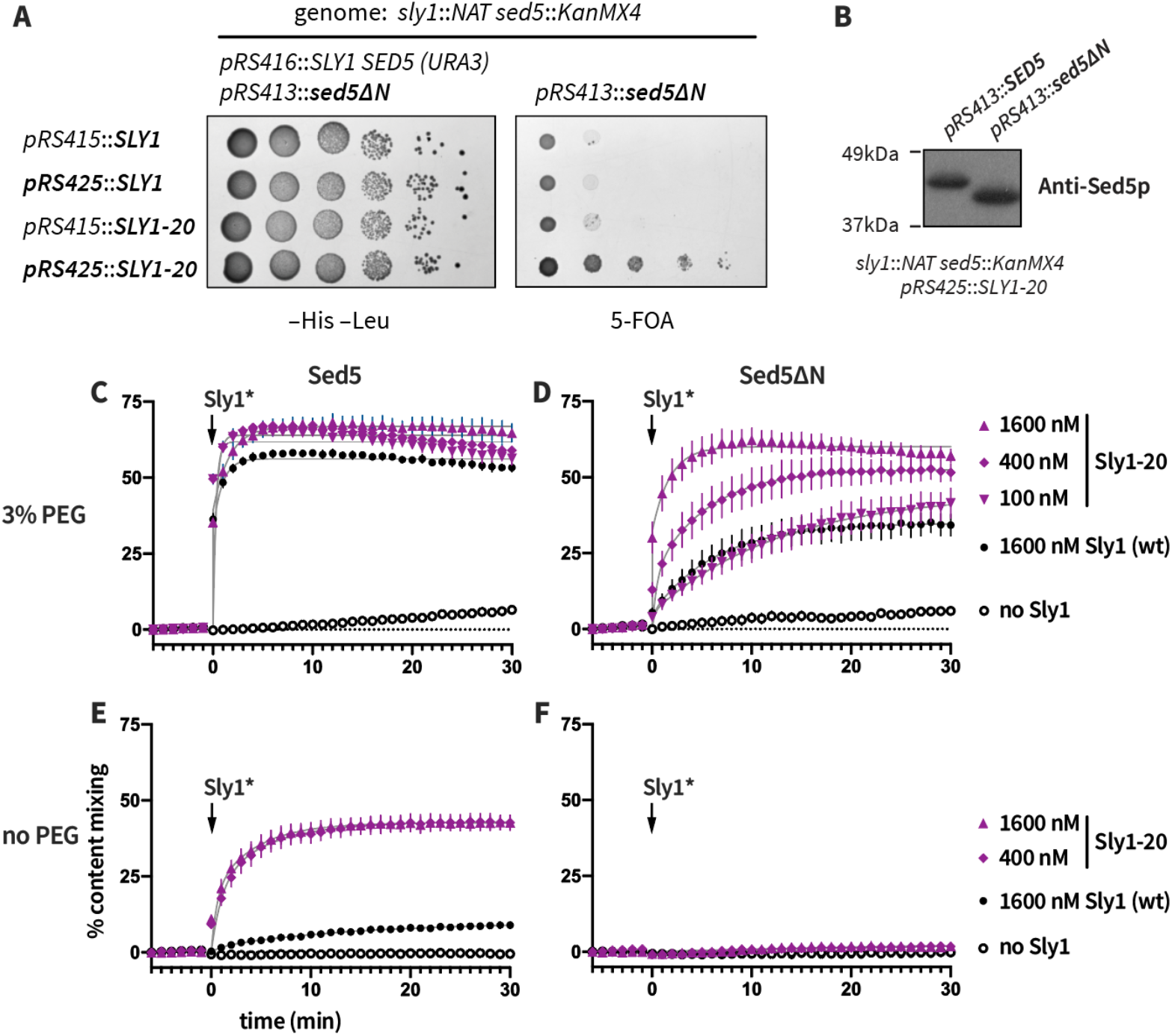
High concentrations of Sly1-20 bypass loss of Sed5 N-peptide or tethering, but not both. **A**, *In vivo* lethality of *sed5ΔN* is suppressed by *SLY1-20* expressed from a multiple-copy plasmid (pRS425) but not from a single-copy plasmid (pRS415). Growth assays were performed as in Fig. 1; a more extensive set of controls is presented in Supplementary Fig. S1. **B**, In the presence of high-copy Sly1-20, the abundance of Sed5ΔN is similar to that of wild type Sed5. Cell extracts were prepared and analyzed by immunoblotting with anti-Sed5 antiserum. **C-F**, Reactions were set up with RPLs, Sec17, Sec18, Mg2+·ATP, and 3% (C,D) or 0% (E,F) PEG. Q-SNARE liposomes bore either wild type Sed5 (C,E) or Sed5ΔN (D,F). The reactions were incubated for 5 min and fusion was initiated by adding Sly1 or Sly1-20 at t = 0. Points show mean ±sem of at least three independent experiments. Gray lines show least-squares fits of a second-order kinetic function.

Genetic suppression can occur through direct or indirect mechanisms. Thus, we used the RPL system to test whether suppression of *sed5ΔN* by multicopy *SLY1-20* occurs through a direct or indirect mechanism. In the presence of PEG, Sly1-20 at 1600 nM was able to drive fusion of Sed5ΔN RPLs to nearly the rates and extents seen with wild type Sed5 RPLs and wild type Sly1 at 100 nM (compare **Figs. 3C** and **D**). However, in the absence of PEG (that is, under tethering-deficient conditions), Sly1-20 was unable to drive fusion of RPLs bearing Sed5ΔN, even at the highest concentrations tested (**Fig. 3E,F**). Taken together, the fusion results closely mirror the *in vivo* matrix of genetic interactions among *SED5* and *SLY1* alleles. When wild-type Sly1 is present, fusion is severely attenuated if the Sed5 N-peptide is deleted, but high concentrations of Sly1-20 can rescue Sed5Δ21. The *in vitro* experiments show that this rescue occurs through direct effects on the fusion machinery. With wild-type Sed5, Sly1-20 at moderate concentrations can compensate for tethering deficiencies either *in vitro* (0% PEG) or *in vivo* (*e.g.*, *ypt1* or *uso1* deficiency). However, consistent with *in vitro* tethering assays (Duan et al., submitted), our fusion experiments show that hyperactive Sly1-20 cannot compensate simultaneously for loss of the Sed5 N-peptide and reduced tethering.

### Sly1 can stimulate fusion independently of Sed5 opening

Sly1 is reported to open closed Sed5, to allow assembly of SNARE core complexes (Demircioglu et al., 2014). We hypothesized that Sed5 opening might be only one of multiple mechanisms through which Sly1 stimulates fusion. To test this idea, we prepared RPLs bearing two different Sed5 mutants that cannot adopt a closed conformation (**Fig. 4A**). Sed5ΔHabc lacks the autoinhibitory Habc domain required to form a closed conformation, but still has the N-terminal 21 amino acids which bind Sly1 with high affinity. Sed5ΔN-Habc lacks both the N-peptide and the Habc domain.

**Fig. 4.**
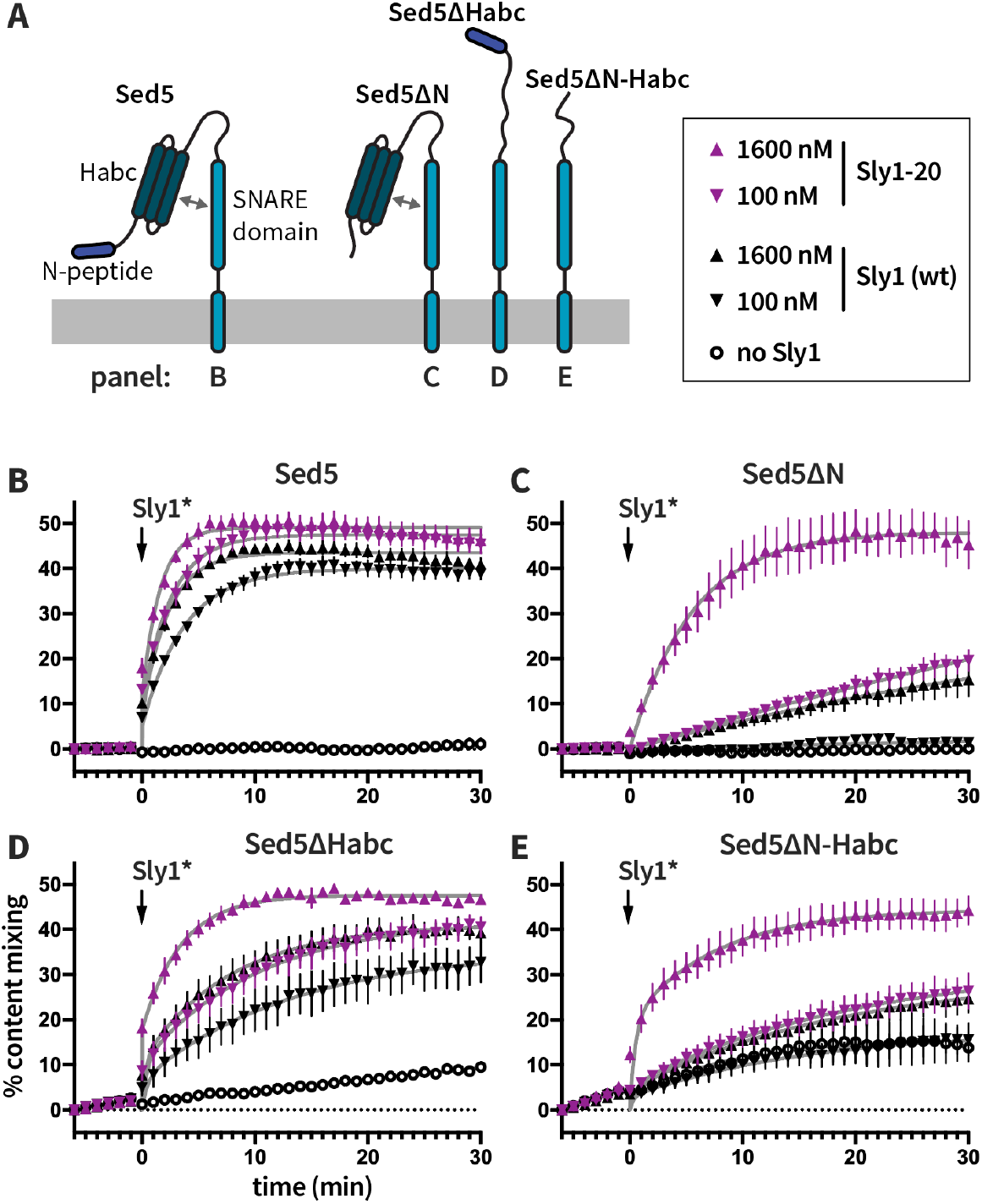
Sly1 stimulates fusion driven by constitutively open Sed5. **A**, Diagram showing Sed5 constructs used in this figure. Sed5ΔN lacks residues 1-21. Sed5 ΔHabc lacks residues 51-180. Sed5ΔN-Habc lacks residues 1-180. **B**. RPLs bearing Sed5 without the Habc domain exhibit slow, Sly1-independent fusion. **D-G**. Sly1 and Sly1-20 stimulation of fusion by RPLs bearing the Sed5 mutants indicated. At t = −6 min., reactions were initiated with the indicated RPLs in the presence of 3% PEG, 100 nM Sec17, 100 nM Sec18, and 1 mM Mg·ATP. At t = 0, Sly1 or Sly1-20 was added to the reactions at 0, 100, or 1600 nM, as indicated in the legend in the upper right corner. Points show mean ±s.e.m. of at least three independent experiments. Gray lines show least-squares fits of a second-order kinetic function.

*In vivo*, both *sed5ΔHabc* and *sed5ΔN-Habc* confer recessive lethal phenotypes. The lethality of these alleles was not suppressed by expression of *SLY1* or *SLY1-20*, from either single- or multiple-copy vectors (**Supplementary Fig. S2A**). Analyses in *SED5*/*sed5* cells indicated that the protein products of the *sed5* mutant alleles are indeed synthesized (**Supplementary Fig. S2B**). However, these mutant proteins are mis-localized and ultimately degraded in the lumen of the lysosomal vacuole (**Supplementary Fig. S2C**). The Habc domain therefore contains information essential for correct Sed5 localization and *in vivo* function. Two of us have recently performed a more detailed analysis of Sed5 localization determinants (Gao and Banfield, 2019).

To assess the role of Sed5 autoinhibition in membrane fusion we returned to the *in vitro* RPL assay system. As before, RPLs bearing wild type Sed5 or Sed5ΔN exhibited little or fusion when Sly1 was absent (**Fig. 4 B,C**; black open circles). In contrast, RPLs bearing either Sed5ΔHabc or Sed5ΔN-Habc exhibited spontaneous but slow Sly1-independent fusion (**Fig. 4D,E**; black open circles). These results strongly corroborate solution biochemistry studies which indicate that autoinhibition by the Sed5 Habc domain prevents SNARE core complex assembly (Demircioglu et al., 2014). In the presence of Sly1 or Sly1-20, however, fusion was stimulated whether the Habc domain was present (**Fig. 4B,C**) or absent (**Fig. 4D,E**). We conclude that although one function of Sly1 is to open the closed conformation of Sed5, Sly1 must also have additional fusion-promoting activities.

### Sly1 can stimulate fusion independently of Sed5 opening and close-range tethering

An auto-inhibitory loop conserved among Sly1 family members harbors a close-range vesicle tethering activity, and this activity is indispensable for the hyperactivity of the Sly1-20 mutant (Duan et al., submitted). We therefore asked whether Sly1 can stimulate fusion independently of *both* its close-range tethering function and its ability to open the Sed5 closed conformation. Reactions were initiated with RPLs bearing Sed5ΔHabc (which cannot adopt a closed conformation), and with wild type Sly1 or Sly1 mutants defective in close-range tethering (**Fig. 5A**). The Sly1Δloop mutant lacks the entire Sly1-specific regulatory loop including the amphipathic helix α21, which is required for close-range membrane tethering. When Sly1Δloop was added to reactions containing Sed5-ΔHabc RPLs, an increase in fusion was still observed, indicating that Sly1 must have fusion-stimulating activities beyond Sed5 opening and close-range tethering.

**Fig. 5.**
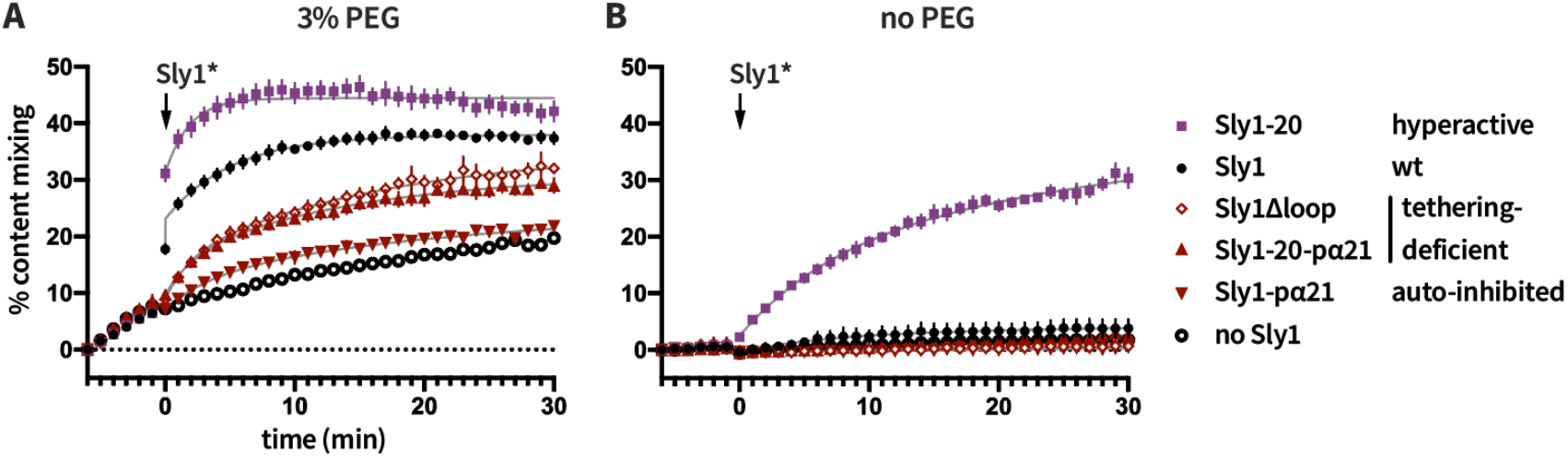
Sly1 can stimulate fusion independently of both Sed5 opening and close-range tethering. At t = −6 min., reactions were initiated with R-SNARE RPLs, Q-SNARE RPLS bearing Sed5ΔHabc, 100 nM Sec17, 100 nM Sec18, and 1 mM Mg·ATP. The reactions also contained (**A**) 3% PEG or (**B**) 0% PEG. At t = 0, the indicated Sly1 variants were added to 1600 nM final, as indicated in the legend. Points show mean ±sem of at least three independent experiments. Gray lines show least-squares fits of a second-order kinetic function.

This conclusion was buttressed by two additional Sly1 mutants. In the Sly1pα21 protein, five apolar residues within helix α21 are mutated, preventing the loop from binding to membranes. Sly1-pα21 has at least two functional defects: it is constitutively auto-inhibited, and it is defective for close-range tethering (Duan et al., submitted). As expected, Sly1-pα21 stimulated only barely detectable fusion above background (**Fig. 5A**). In the compound mutant Sly1-20-pα21, autoinhibition is released (the loop is open), but close-range tethering is still compromised. When added to reactions with Sed5ΔHabc RPLs, Sly1-20-pα21 stimulated fusion similarly to Sly1Δloop (**Fig. 5A**). When the same panel of Sly1 variants was tested with Sed5ΔHabc RPLs under tethering-deficient conditions (**Fig. 5B**), only Sly1-20 (constitutively open and presenting helix α21) was able to stimulate substantial fusion. Thus, fusion of Sed5ΔHabc RPLs requires a tethering activity which can be provided either by the tethering-hyperactive Sly1-20, or by PEG. We conclude that both Sly1 opening of closed Sed5, and Sly1 close-range tethering activity, contribute to the ability of Sly1 to promote fusion, and that Sly1 has one or more additional fusion-promoting activities. In Vps33 and Munc18-1, domain 3a appears to serve as a template for *trans*-SNARE complex assembly (Baker et al., 2015; Jiao et al., 2018). The near-inability of the constitutively auto-inhibited mutant Sly1-pα21 to stimulate fusion (**Fig. 5A**) implies that the additional Sly1 activities involve Sly1 domain 3a, which is occluded when Sly1 is auto-inhibited (Baker et al., 2015; Bracher and Weissenhorn, 2002).

### Close-range tethering by Sly1 promotes assembly of trans-SNARE complexes

Assembly of *cis*-SNARE complexes occurs spontaneously and competes with assembly of fusion-active *trans*-complexes. To test the hypothesis that close-range tethering by Sly1 specifically favors *trans*-SNARE complex assembly, we evaluated the inhibitory activity of the soluble R-SNARE domain of Sec22 (Sec22_SN_-GFP; **Fig. 6**). Reactions were initiated with RPLs bearing the three Q-SNARES along with Sec22_SN_-GFP (0, 2 or 20 μM), and wild type or mutant forms of Sly1. The reactions were incubated for 15 min, then R-SNARE RPLs were added and the reactions were incubated for an additional 6 min. To initiate fusion PEG was added (*t* = 0). PEG was added to 4% final rather than 3% (as in previous experiments) in an effort to compensate for the tethering defects of the Sly1-20-α21p and Sly1Δloop mutants. Reactions driven by wild-type Sly1 or hyperactive Sly1-20 were partially inhibited by 2 μM Sec22_SN_-GFP (**Fig. 6B**), and almost totally inhibited by 20 μM Sec22_SN_-GFP (**Fig. 6C**). In contrast, reactions containing Sly1-20-α21p or Sly1Δloop were almost totally inhibited at both 2 and 20 μM Sec22_SN_-GFP (**Fig. 6B,C**). This indicates that reactions driven by tethering-deficient Sly1 mutants are more sensitive to inhibition by *cis*-complex assembly.

**Fig. 6.**
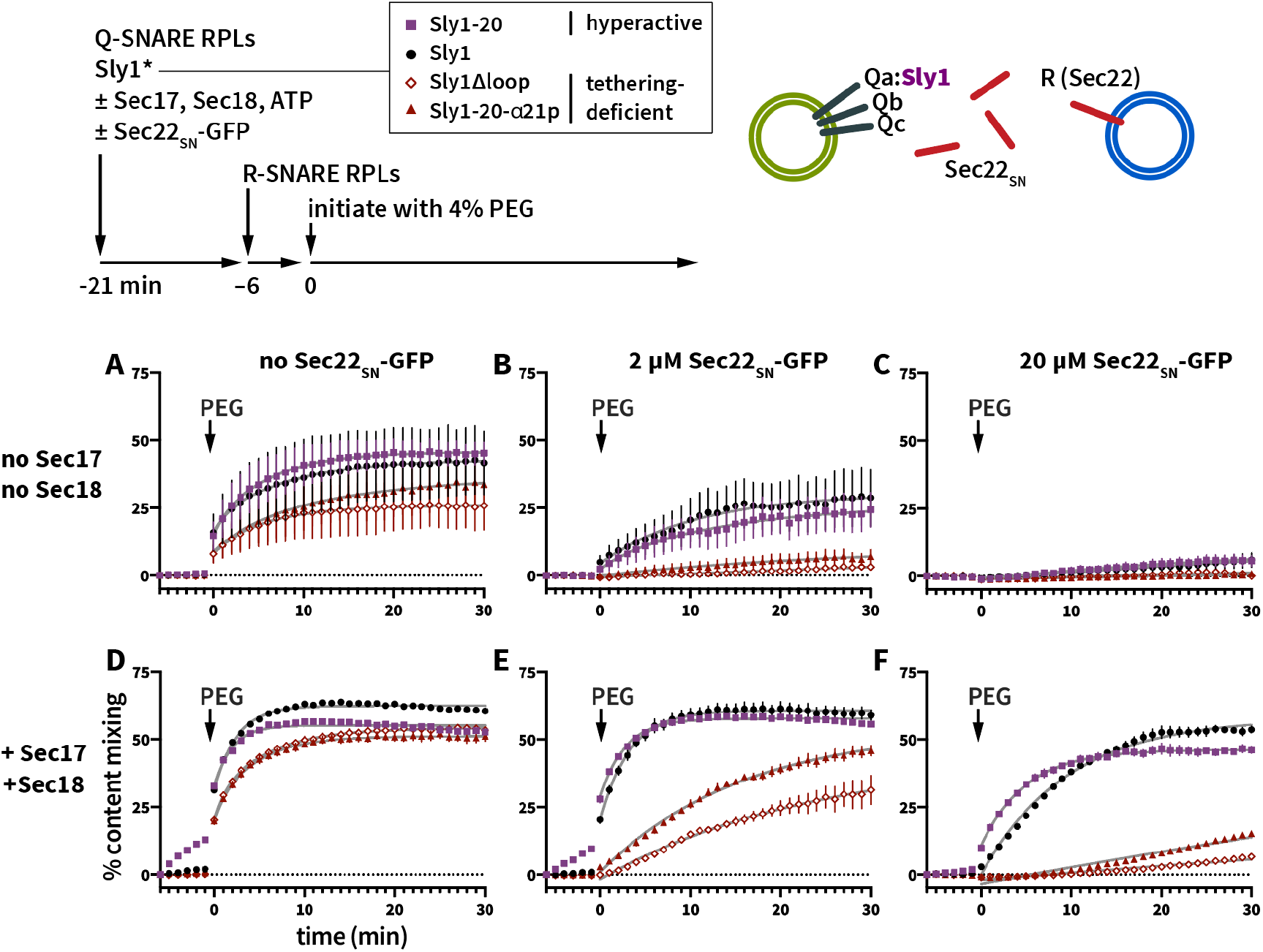
Helix α21 promotes selective formation of fusion-active *trans*-SNARE complexes. At t = −21 min., Q-SNARE RPLs bearing Sed5-WT were mixed with 100 nM SLY1 variants as indicated in the legend, and either without (**A-C**) or with (**D-F**) Sec17, Sec18 (100 nM each) and Mg·ATP (1 mM), and with either 0 μM (**A,D**), 2 μM (**B,E**) or 20 μM (**C,F**) soluble Sec22_SN_-GFP. R-SNARE RPLs were added at t = −6 min. At t = 0, the reactions were initiated by addition of 4% PEG. Points show mean ±sem of at least three independent experiments. Gray lines show least-squares fits of a second-order kinetic function.

We next tested whether inhibition of fusion by Sec22_SN_-GFP can be reversed by driving cycles of SNARE complex disassembly, to generate free SNAREs on the RPLs. Reactions were initiated as in **Fig. 6A-C**, but with Sec17, Sec18, and Mg^2+^·ATP (**Fig. 6D-F**). Reactions containing Sly1 or hyperactive Sly1-20 exhibited efficient fusion in the presence of Sec17/18, even at 20μM Sec22_SN_-GFP. In contrast, reactions driven by the tethering deficient Sly1 mutants exhibited far less rescue. Similar results were obtained when reactions were run under the same conditions, but with 3% rather than 4% PEG (**Supplementary Fig. S3**). Taken together these experiments support the hypothesis that α21-mediated tethering favors assembly of fusogenic *trans*-SNARE complexes versus non-fusogenic *cis*-SNARE complexes, even when the pool of unpaired SNAREs is continuously replenished through cycles of Sec17- and Sec18-mediated *cis*-SNARE complex disassembly.

We wondered whether similar results might be obtained under more physiologically relevant conditions, using wild-type membrane-anchored Sec22 rather than soluble Sec22. We therefore assayed the fusion of 4-SNARE QabcR RPLs with 3-SNARE Qabc RPLs, in the presence of wild type and mutant Sly1 variants (**Fig. 7**). Because QabcR RPLs can form quaternary SNARE complexes *in cis* as well as *in trans*, they require Sec17/18 for efficient fusion. In this configuration the formation of *trans*-SNARE complexes and the re-formation of *cis*-SNARE complexes are competing processes. As above, we ran reactions in the presence of 4% PEG in an effort to compensate for the defects of tethering-deficient Sly1 mutants. Even given the results with soluble Sec22, we were surprised at the clarity of the results: Sly1-20-α21p and Sly1Δloop were almost fusion-inactive when confronted with the QabcR SNARE topology (**Fig. 7A**), even when the Sly1 mutants were supplied at 1600 nM (**Fig. 7D**) and despite the presence of 4% PEG. These defects are considerably more severe than the defects observed in RPL experiments using the Qabc vs. R-SNARE topology, where QabcR *cis*-SNARE complexes cannot form during the first round of fusion (e.g., Fig. 6a; and Duan et al., submitted, Fig. 5B,D). Two mutants hyperactive for close-range tethering, Sly1-20 and Sly1-T559I (Duan et al., submitted), exhibited activity as strong as or stronger than the wild type.

**Fig. 7.**
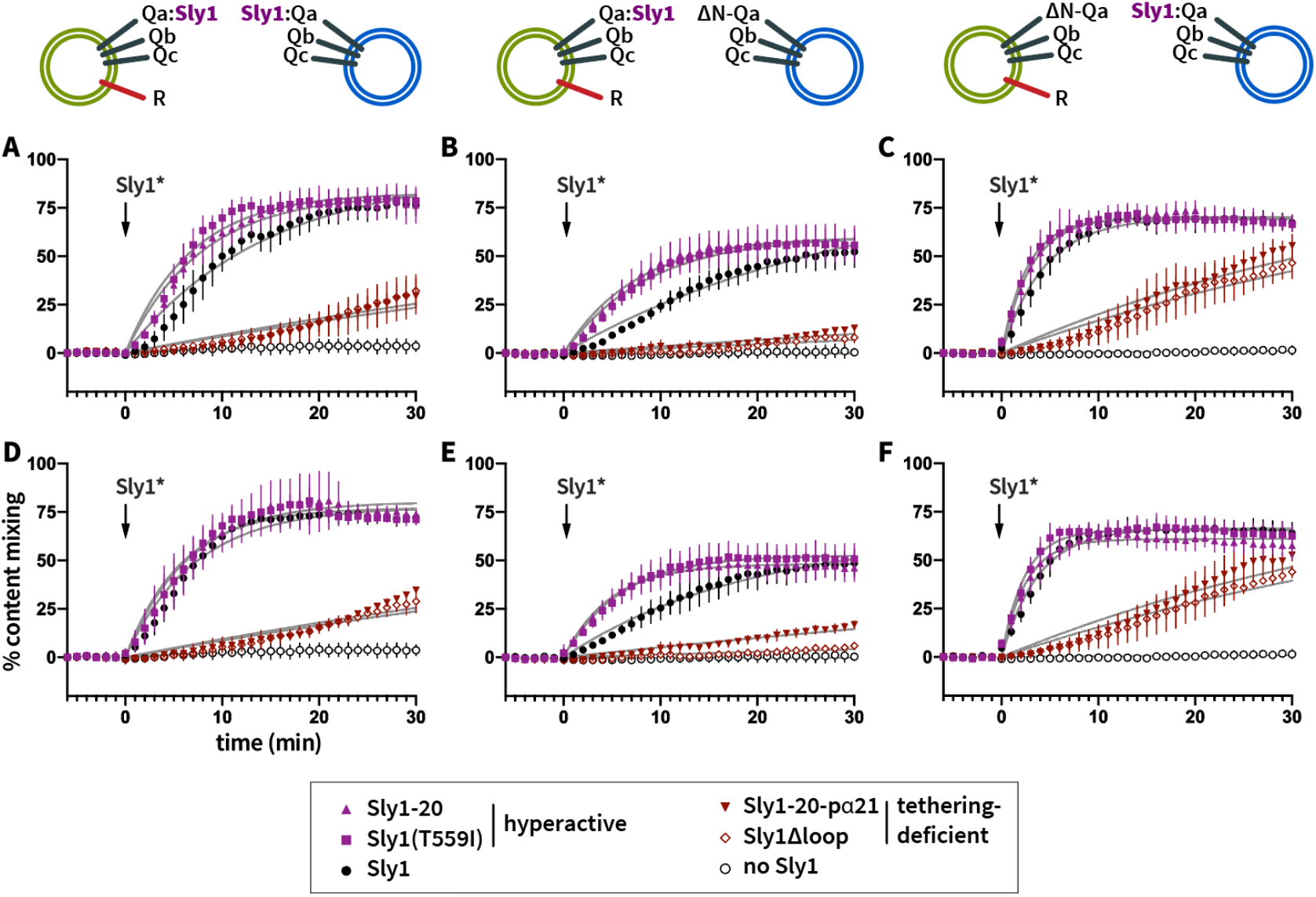
Helix α21 promotes Sly1 discrimination between *cis*- and *trans*-SNARE complexes. Reactions were initiated with RPLs bearing SNAREs in the indicated topologies. All reactions contained Sec17, Sec18 (100 nM each), and Mg^2+^·ATP (1 mM). PEG was added to 4% final rather than 3% to assist the tethering-deficient Sly1 mutants. Fusion was initiated (t = 0) by adding the indicated Sly1 mutants to either 100 nM (**A-C**) or 1600 nM (**D-F**). Points show mean ±sem of at least three independent experiments. Gray lines show least-squares fits of a second-order kinetic function.

As Sly1 is recruited to the Qa-SNARE Sed5 through a high-affinity interaction with the Sed5 N-peptide, we used the Sed5ΔN mutant protein to direct Sly1 primarily to either the QabcR RPLs (**Fig. 7B,E**), or to the Qabc RPLs (**Fig. 7C,F**). The results show that fusion occurs most rapidly when Sly1 is placed *in trans* to the R-SNARE, Sec22 (**Fig. 7C,F**) — the configuration where R-SNARE binding *in trans* is most likely and where *cis* QabcR complex assembly is least likely to be stimulated by Sly1. Fusion is slowest when Sly1 is placed *in cis* to the R-SNARE; under this condition, the tethering-deficient Sly1 mutants are almost totally unable to drive fusion (**Fig. 7C,D**). Taken together the experiments in Figs. 6 and 7 demonstrate that the Sly1 close-range tethering activity takes on special importance when Sly1 must selectively catalyze formation of *trans*-SNARE complexes in competition with formation of inactive *cis*-SNARE complexes.

### Soluble Sed5 Habc domain promotes Sly1-stimulated fusion in vitro

In our experiments comparing various Sed5 mutants, we were surprised to see that although Sed5ΔHabc can catalyze spontaneous Sly1-independent fusion, its ability to support Sly1-stimulated fusion was reduced compared to wild type Sed5 (compare **Figs. 4B** and **4D**). These results suggested that the Sed5 Habc domain, in addition to being autoinhibitory, might have a positive, fusion-promoting activity. To test this hypothesis we asked whether the Sed5 Habc domain, supplied in soluble form, would alter the ability of Sly1 and its cognate SNAREs to drive fusion.

Reactions were initiated with Qabc-SNARE RPLs bearing four different Sed5 variants: wild-type, Sed5ΔN, Sed5ΔHabc or Sed5 ΔN-Habc. Each of these reactions was performed in the absence or presence of soluble Sed5 Habc or N-Habc (**Fig. 8A-D**). Because Sly1 binds to soluble N-Habc domain with sub-nanomolar affinity (Demircioglu et al., 2014), Sly1 and soluble Habc or N-Habc were pre-mixed before they being added together to the reaction mixture. The reactions were initiated without PEG and monitored for 6 min. To initiate fusion, PEG was added to 3% (t = 0). To our surprise, both the Habc and N-Habc domains of Sed5 stimulated fusion in a dose-dependent manner. Fusion was most efficiently stimulated when the N-peptide was present on the soluble Habc domain (**Fig. 8B**), rather than on the membrane-bound Sed5 (**Fig. 8C**), and less efficiently stimulated when N-peptides were present on both Sed5 and the soluble N-Habc protein (**Fig. 8A**). However, Habc stimulated fusion under every condition tested, even when the N-peptide was absent from both Sed5 and the soluble Habc domain (**Fig. 8D**).

**Fig. 8.**
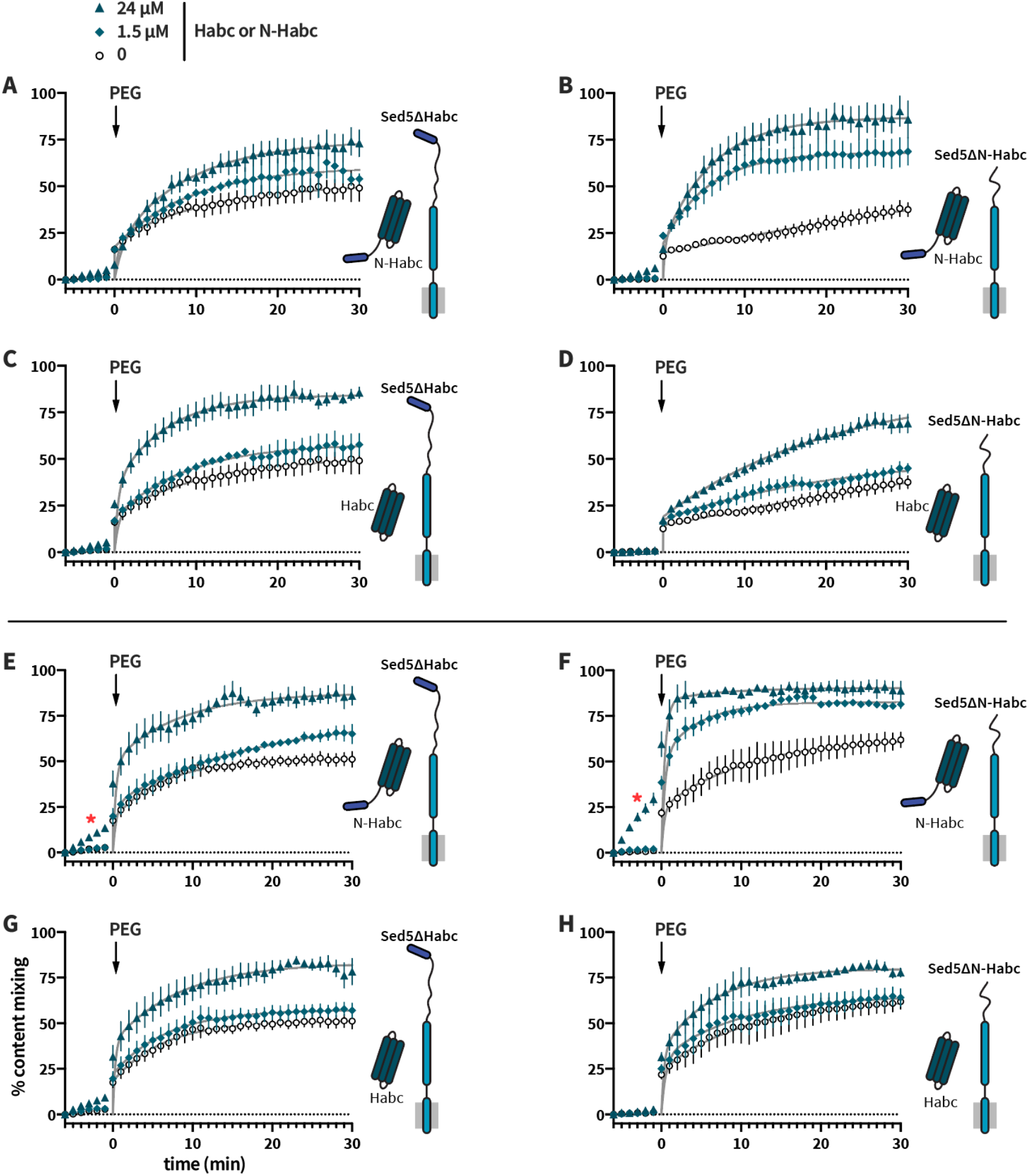
Soluble Sed5 Habc domain can stimulate Sly1-dependent fusion. At t = −6 min., R-SNARE RPLs and Q-SNARE RPLS bearing either Sed5ΔHabc or Sed5ΔN-Habc were mixed either without (**A-D**) or with (**E-H**) Sec17, Sec18 (100 nM), and Mg^2+^·ATP (1 mM). Reactions contained 100 nM Sly1 that had been preincubated with either soluble Sed5N-Habc (**A,B,E,F**) or Sed5Habc (**C,D,G,H**). At t = 0, the reactions were initiated by addition of 3% PEG. Points show mean ±sem of at least three independent experiments. Gray lines show least-squares fits of a second-order kinetic function. In **E** and **F**, red asterisks (*) indicate fusion that occurred prior to addition of PEG.

Similar results were obtained in reactions lacking (**Fig. 8A-D**) or containing Sec17 and Sec18 (**Fig. 8E-H**). Remarkably, in the presence of Sec17 and Sec18 and at the highest concentration of N-Habc, robust fusion was observed even before PEG was added to the reaction (**Fig. 8E,F**; red asterisks). This indicates bypass of the tethering requirement — a phenotype previously observed only with hyperactive Sly1 mutants such as Sly1-20 or at very high concentrations of PEG (Duan et al., submitted). We conclude that the Sed5 Habc domain augments the efficiency of Sly1-stimulated fusion, that the Habc domain need not be covalently coupled to Sed5. In other words, “split Sed5” can drive fusion.

### Split Sed5 can function *in vivo*

Next, we tested whether soluble Sed5 Habc or N-Habc domains might suppress the lethal phenotype of cells expressing Sed5 variants lacking the Habc domain (**Fig. 9**). Single-copy dicistronic plamids were constructed bearing *sed5ΔN-Habc* or *sed5ΔHabc*, as well as either *Habc* or *N-Habc*. These test plasmids were introduced into *sly1Δ sed5Δ* strains harboring a counter-selectable *SLY1 SED5* balancer plasmid. *In vitro*, we had noted that Qabc RPLs bearing wild-type Sed5 fuse with similar efficiency when either Sly1 or hyperactive Sly1-20 are supplied, but that RPLs bearing Sed5ΔHabc are considerably more responsive to Sly1-20 (**Fig. 4**). Thus, we also tested the effects of single-copy or multiple-copy plasmids bearing either *SLY1* or *SLY1-20*. The results show that the Sed5 N-Habc domain can support viability when present solely as a soluble fragment (**Figs 9A,B**). However, viability of these “split Sed5” cells requires Sly1 hyperactivity. Only *SLY1-20* expressed from a high-copy vector supported robust growth with split Sed5. Single-copy *SLY1-20* facilitated very slow growth. Moreover, as in the *in vitro* assays, rescue was most robust when the N-peptide is on the soluble Habc fragment (**Figs9 A,B**). No rescue was observed in cells totally lacking the Sed5 N-peptide (**Fig. 9C**) and rescue was only barely detectable when the N-peptide was present solely on the membrane-anchored mutant Sed5 (**Fig. 9D**).

**Fig. 9.**
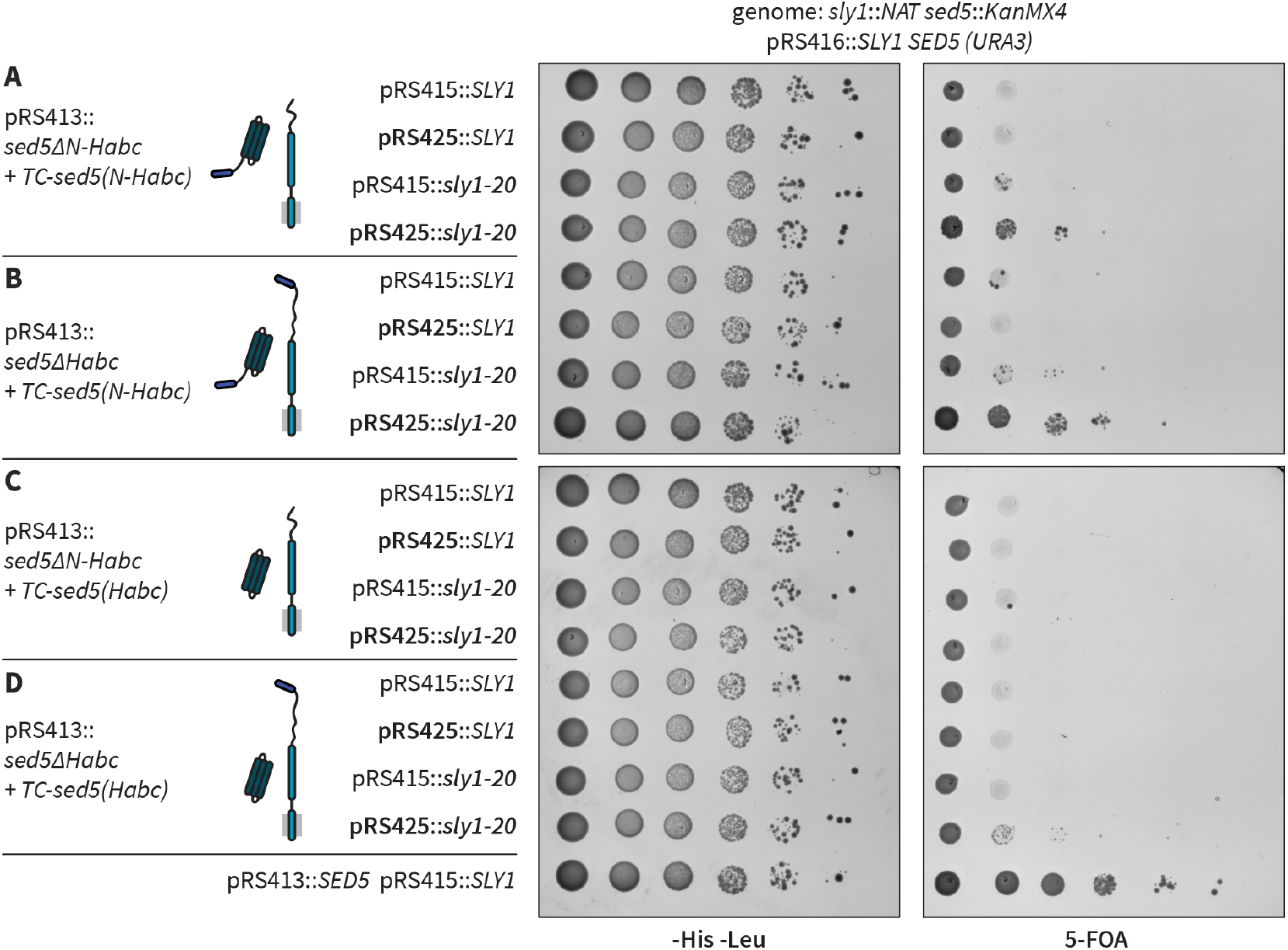
Soluble Sed5 N-Habc fragment supports viability of cells in the presence of *SLY1-20*. Cells with the indicated genotypes were constructed as described in the text. These strains were grown in liquid media -His -Leu media to maintain the plasmids. Serial dilutions were then plated to either -His -Leu solid media or to solid media containing 5-FOA, to eject the *SLY1 SED5* balancer plasmid. Expression of the membrane-anchored Sed5 variants was driven using the native *SED5* promoter and terminator. Expression of the soluble Habc and N-Habc fragments was driven using the strong *TPI1* promoter and the *<u>C</u>YC1* terminator (*TC*). Immunoblot analyses of Sed5* expression in these cells are shown in Supplementary Fig. S4.

Immunoblot analyses of whole cell lysates from cells expressing wild-type Sed5 indicated that the the truncated Sed5 variants and soluble fragments were expressed (**Supplementary Fig. S4A**). In cells lacking the *SLY1 SED5* balancer plasmid and expressing either Sed5ΔN-Habc or Sed5ΔHabc, the steady-state level of the soluble N-Habc fragment depended on the gene dosage of *SLY1-20*, suggesting that the stability of the soluble fragment is controlled by its interaction with Sly1-20 protein (**Supplementary Fig. S4B**). Taken together, the *in vitro* and *in vivo* results here and in (Gao and Banfield, 2019) indicate that, in addition to being auto-inhibitory, the Sed5 Habc domain has positive functions: it both promotes Sly1-dependent membrane fusion, and regulates Sed5 subcellular localization.

## DISCUSSION

Our experiments show that Sly1 has multiple distinguishable activities which promote SNARE-mediated fusion (Fig. 9). First, Sed5-bound Sly1 has the intrinsic ability to tether incoming vesicles through the amphipathic helix α21 within the Sly1 regulatory loop (Duan et al., submitted). Second, Sly1 has the ability to open the closed, auto-inhibited conformation of the Qa-SNARE Sed5 (Demircioglu et al., 2014). When these activities are experimentally bypassed, Sly1 still promotes fusion (albeit less efficiently) through a third activity that probably involves domain 3a. We infer that the third function is probably the selective and accurate nucleation of *trans*-SNARE complex assembly. These functions are interlinked. Defects in the Sly1 close-range tethering function result in dramatically impaired fusion when *cis*-SNARE and *trans*-SNARE complex assembly are competing processes, indicating that the tethering function is closely coupled to selective catalysis of *trans*-SNARE complex assembly. Although the precise mechanism through which this occurs is not yet clear, structural data are suggestive. If the R-SNARE Sec22 binds to Sly1 in a configuration similar to the binding of the R-SNARE Nyv1 to Vps33 (Baker et al., 2015), then the open Sly1 loop should tether the incoming vesicle in an orientation optimal for capture of the vesicular R-SNARE. We therefore speculate that the close-range tethering mechanism serves not only to inspect incoming vesicle membranes and to trigger Sly1 activation, but to steer Sly1 into a spatial orientation that maximizes the likelihood of productive R-SNARE binding to domain 3a. When *cis*-SNARE complexes can form in competition with *trans*-complexes, tethering-defective Sly1 mutants exhibit profound fusion defects, even when tethering is stimulated by 4% PEG (Figs. 6 and 7).

**Fig 9.**
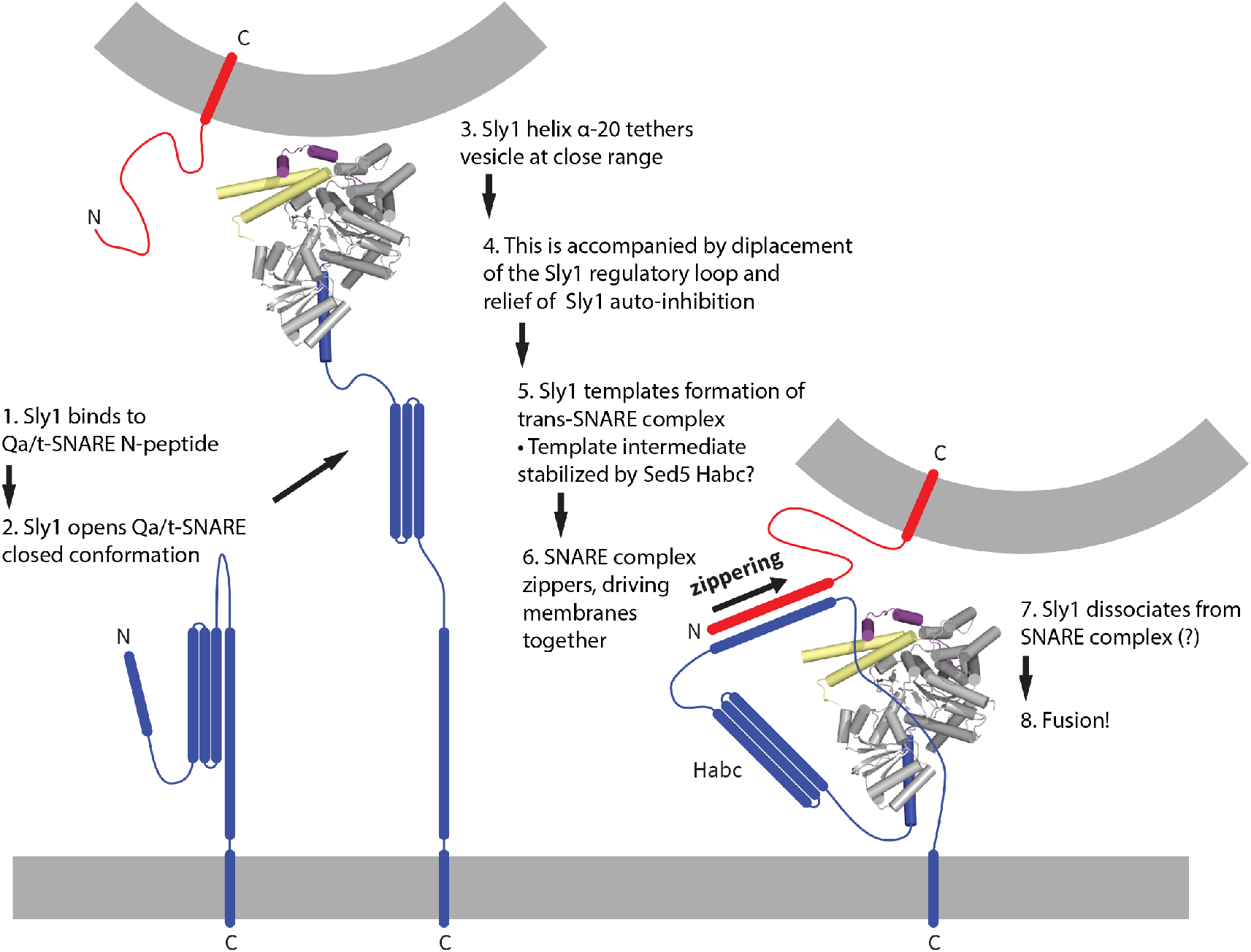
Working model.

At the presynaptic nerve terminal the SM protein UNC-18/Munc18-1 seems to lock its Qa-SNARE, Syntaxin-1A, into a closed conformation. Another protein, UNC-13/Munc13-1 appears to be primarily responsible for opening syntaxin-1A (Richmond et al., 2001; Wang et al., 2017; Yang et al., 2015). However, it is unclear whether proteins homologous or analogous to UNC-13 are required for Qa-SNARE opening during other SNARE-mediated fusion events (Pei et al., 2009). Like Syntaxin-1A, the ER-Golgi Qa-SNARE Sed5 is normally autoinhibited. In contrast to Munc-18, Sly1 opens the closed Qa-SNARE (Demircioglu et al., 2014). However, Sly1 still stimulates function of Sed5ΔHabc mutants that cannot close. This was, perhaps, expected. Although all Qa (syntaxin-family) SNARE proteins have trihelical Habc domains, Habc domains are not always auto-inhibitory – yet SM proteins are still needed. For example Vam3, the Qa-SNARE of the yeast lysosomal vacuole, is constitutively open and its Habc domain lacks a groove that might bind to its SNARE domain, yet the SM protein Vps33 is still essential for fusion (Baker et al., 2015; Dulubova et al., 2001; Lobingier et al., 2014; Rieder and Emr, 1997; Seals et al., 2000). Thus, clamping the Qa-SNARE in a closed conformation, and opening the closed conformation, are pathway-specific elaborations rather than activities common to all SMs.

In addition to the above functions Sly1 has additional activities. Sly1 reduces the rate of SNARE complex disassembly by Sec17 and Sec18 (Lobingier et al., 2014). This is consistent with studies of Vps33 and Munc18-1, showing that these SMs protect assembled *trans*-SNARE complexes from premature disassembly by Sec17 and Sec18 (Lobingier et al., 2014; Prinslow et al., 2019; Schwartz et al., 2017; Song et al., 2017; Stepien et al., 2019; Xu et al., 2010; Duan et al., submitted). Thus we reiterate our previous suggestions (Lobingier et al., 2014; Schwartz et al., 2017) that SM proteins are, *sensu stricto*, enzymes.

Like all enzymes, SMs bind substrates (vesicular and target SNARE domains), placing them in a stereoselective orientation that reduces the kinetic barrier to formation of product (the *trans*-SNARE complex); the SMs then dissociate from the product to engage in additional cycles of catalysis (Jiao et al., 2018; Schwartz et al., 2017). Also as expected for true enzymes, SMs prevent off-pathway reactions (*e.g.*, assembly of non-cognate SNARE complexes, or of *cis*-rather than *trans*-complexes). They achieve this increase in specificity through kinetic partitioning within the forward assembly pathway (Hardy and Randall, 1991; Lai et al., 2017; Lambright et al., 1994; Peng and Gallwitz, 2002), and through kinetic proofreading of incorrect SNARE assemblies, since SMs selectively protect cognate SNARE complexes from premature disassembly by proofreading enzymes, while non-cognate complexes are efficiently disassembled (Choi et al., 2018; Lobingier et al., 2014; Prinslow et al., 2019; Schwartz et al., 2017; Song et al., 2017; Xu et al., 2010).

We were somewhat surprised to find that Sly1-dependent fusion driven by Sed5ΔHabc is slower than fusion driven by wild-type Sed5. Similarly, deletion of the Vam3 Habc domain causes a kinetic defect in homotypic vacuole fusion (Laage and Ungermann, 2001; Lürick et al., 2015; Pieren et al., 2010); but see (Wang et al., 2001). When we added soluble Habc domain to reactions containing Sed5ΔHabc RPLs, fusion activity was restored to wild-type or nearly wild-type levels (Fig. 8). The Habc domain of Sed5 therefore must have a positive function. What could this function be? Pieren et al. (2010) suggested that the Vam3 Habc domain might, through an interaction with Vps33, facilitate a transition from lipid mixing to content mixing. However, we have detected no signals consistent with the hypothesis that the Sed5 Habc domain influences the transition from lipid to content mixing. Experiments from the Zhang laboratory are more suggestive of an underlying mechanism. Using single-molecule force spectroscopy, they probed the formation and stability of template complexes consisting of neuronal SNAREs and the cognate SM Munc18-1. Formation of the SNARE–Munc18-1 template complex was almost an order of magnitude less efficient when Syntaxin lacked its N-terminal regulatory domain (the N-peptide and Habc domain). In a striking parallel to our fusion experiments (Figs. 8 and 9), addition of the soluble N-Habc domain rescued template complex formation and increased stability of the template complex by almost an order of magnitude (Jiao et al., 2018). The underlying structural basis for this stabilization is not yet understood.

## MATERIALS & METHODS

### Yeast strains, genetic tests, and microscopy

Yeast and *E. coli* strains are listed in Supplementary Table I. Viability assays were performed as described (Gao and Banfield, 2019).

### Proteins

Full length SNAREs were expressed and purified as described in the companion manuscript. Constructs used to express mutant forms of Sed5 are listed in Supplementary Table 1. Sed5 mutants bearing transmembrane domains were expressed and purified as for full-length wild type Sed5. Soluble domains of Sed5 were expressed and purified as in the companion manuscript. Sly1 and its mutants were expressed and purified as described (Duan et al., submitted).

### RPLs and Fusion assays

RPL lipid compositions, detailed methods for RPL preparation, and the *in vitro* fusion assay are described in the companion study (Duan et al., submitted). In this study additional SNARE topologies were tested, as described in the Results. The molar protein:phospholipid ratio for 4-SNARE liposomes was 1:1200, and 1:600 for Qabc and R SNARE RPLs. For certain experiments, as specified, reactions were set up with a non-standard order of reagent addition, and/or fusion was initiated by adding PEG, rather than by adding Sly1 or its mutants.

## ACKNOWLEDGEMENTS

We are grateful to Drs. R. Baker, I. Topalidou, and M. Zick for helpful advice and critical comments on the manuscript; and D. Beacham (Molecular Probes/Thermo Fisher) for gifts of fluorescent reagents. This work was funded by NIH/NIGMS R01 GM077349 and by University of Washington (AM), and the Hong Kong Research Council GRF 16101718 and AoE/M-05/12-2 (DB)

## SUPPLEMENTARY MATERIALS

**Supplementary Table S1.**
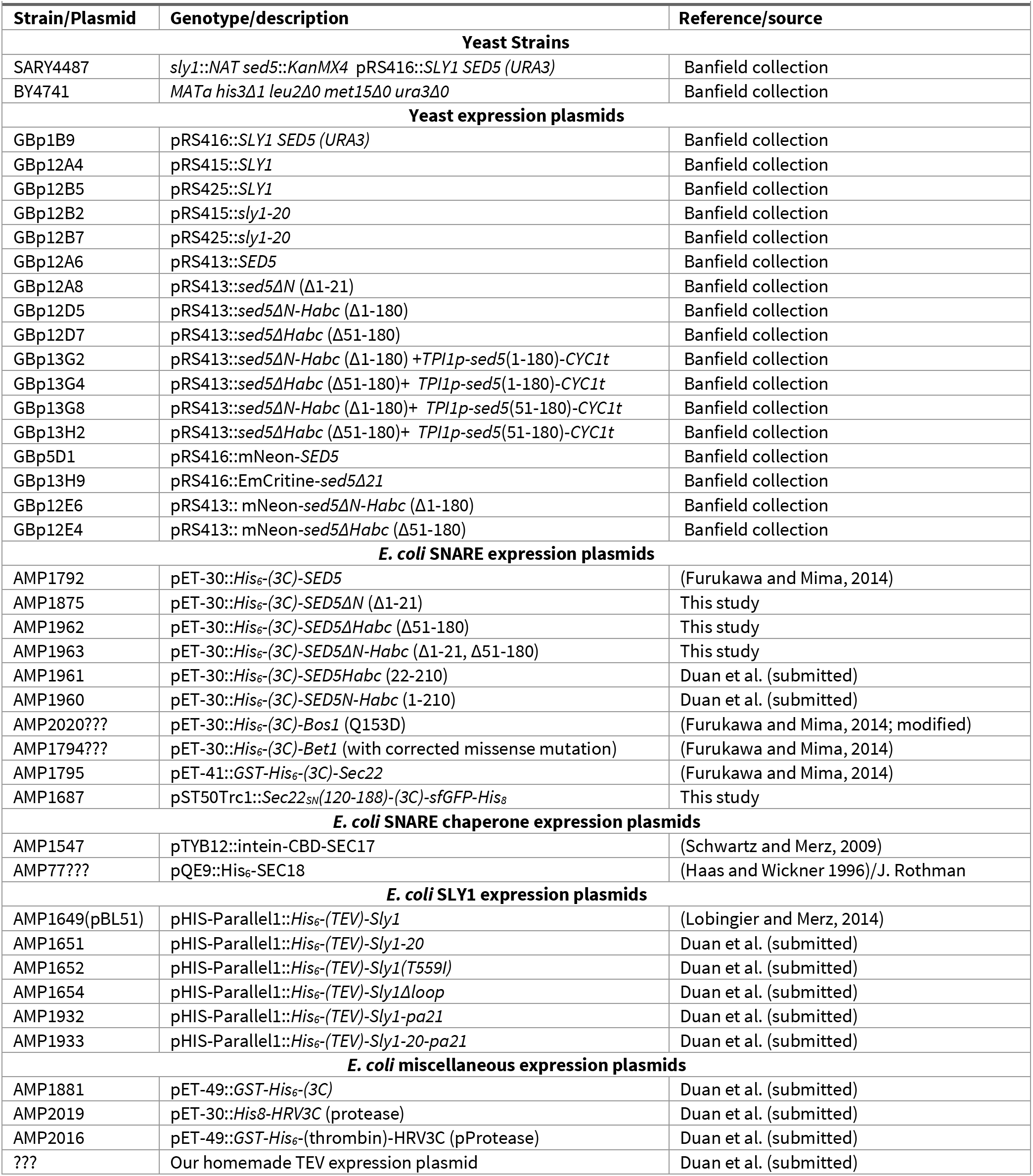
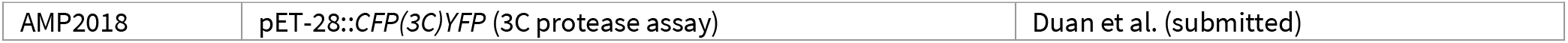
Yeast strains and plasmids used in this study.

**Supplementary Fig. S1.**
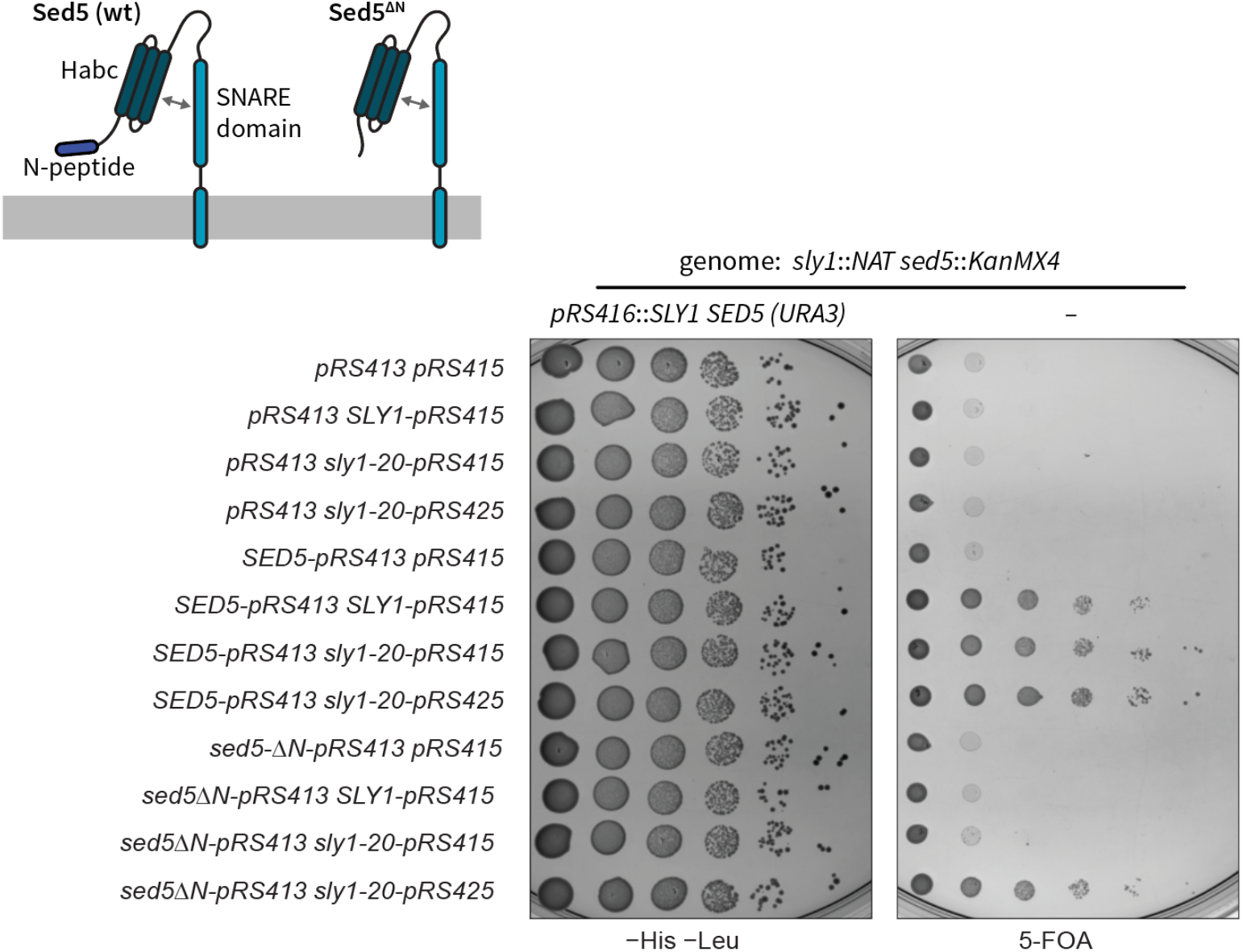
Genetic interactions among *sed5-ΔN*,*SLY1*, and *SLY1-20*. This figure shows the results from Figs. 1B and 3A together, along with additional controls. The *sed5-ΔN* allele was tested using *sed5Δ sly1Δ* double knockout cells that carry intact copies of both *SED5* and *SLY1* on a single counter-selectable plasmid. Forced ejection of the *SED5 SLY1* plasmid by plating onto 5-fluoroorotic acid (5-FOA) resulted in lethality unless both *SED5* and *SLY1* (or *SLY1-20*) were supplied *in trans*.

**Supplementary Fig. S2.**
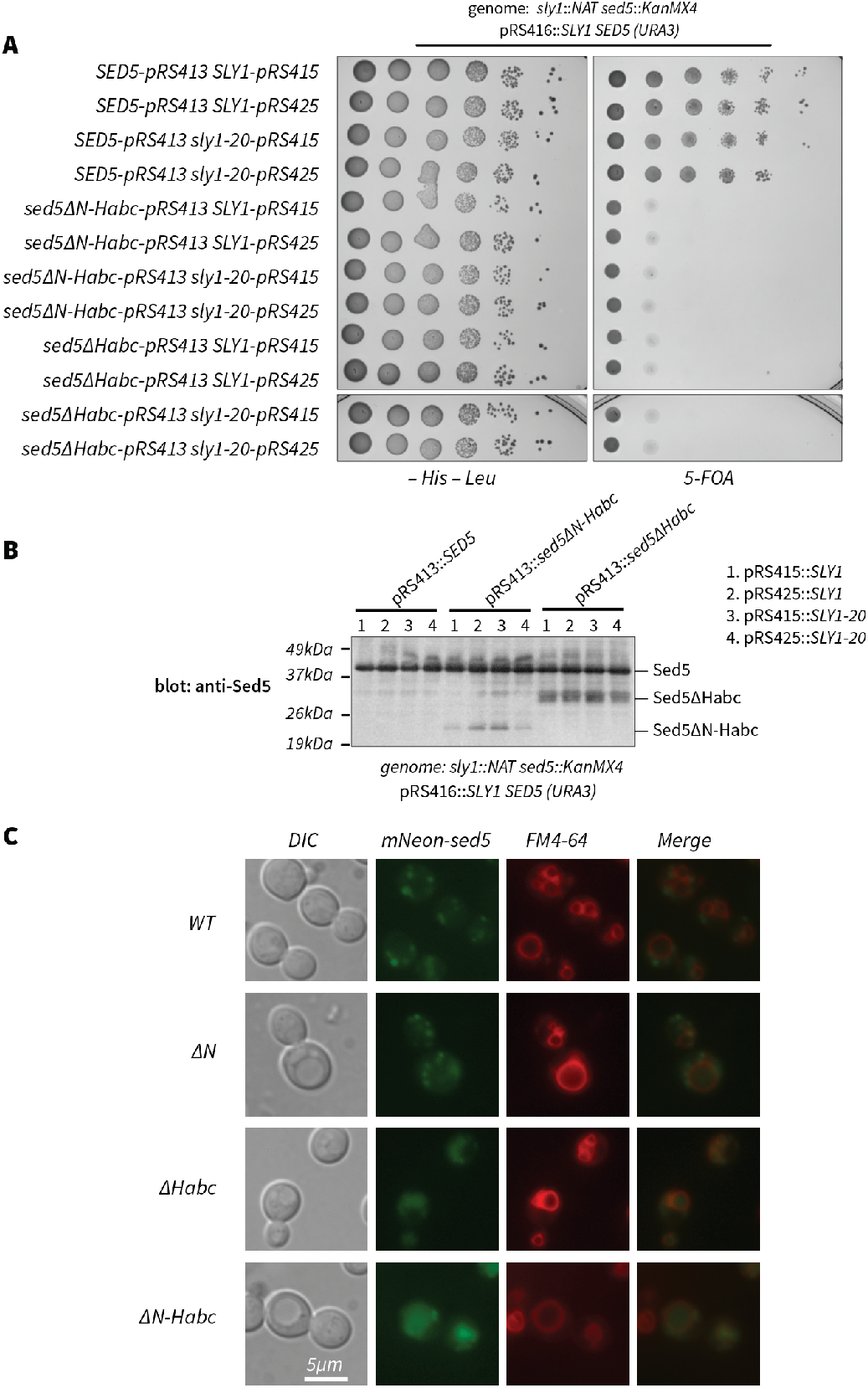
The Sed5 Habc domain is essential *in vivo* and necessary for correct Sed5 localization. **A**, *sed5* alleles encoding variants lacking the Habc domain do not support viability, even in the presence of high-copy *SLY1-20*. **B**, In cells expressing wild type Sed5, Sed5ΔHabc and Sed5ΔN-Habc variants are produced *in vivo* and migrate at the expected sizes. The strains shown in panel A were grown in -His -Leu media. Whole cell lysates were prepared, fractionated on SDS-PAGE, and analyzed by immunoblot with anti-Sed5. Note that the anti-Sed5 antibody is polyclonal. Consequently, band intensities for a given Sed5 variant can be used to infer relative abundance. However, the band intensities cannot be used to infer the relative abundance of different Sed5 constructs. **C**, Subcellular localization of Sed5* variants. Cells expressing both wild-type Sed5 and the indicated mNeon-Sed5* variants were labeled with the vital dye FM4-64 (which marks the vacuolar lysosome), then examined using Nomarski differential interference contrast (DIC) and epifluorescence. Wild-type Sed5 and Sed5ΔN exhibited a punctate localization not overlapping with FM4-64. This is consistent with Golgi localization at steady state. In contrast, Sed5ΔHabc and Sed5ΔN-Habc co-localized with FM4-64, consistent with a pre-vacuolar or vacuolar localization, and appeared to be in the vacuole lumen rather than on the vacuole limiting membrane.

**Supplementary Fig. S3.**
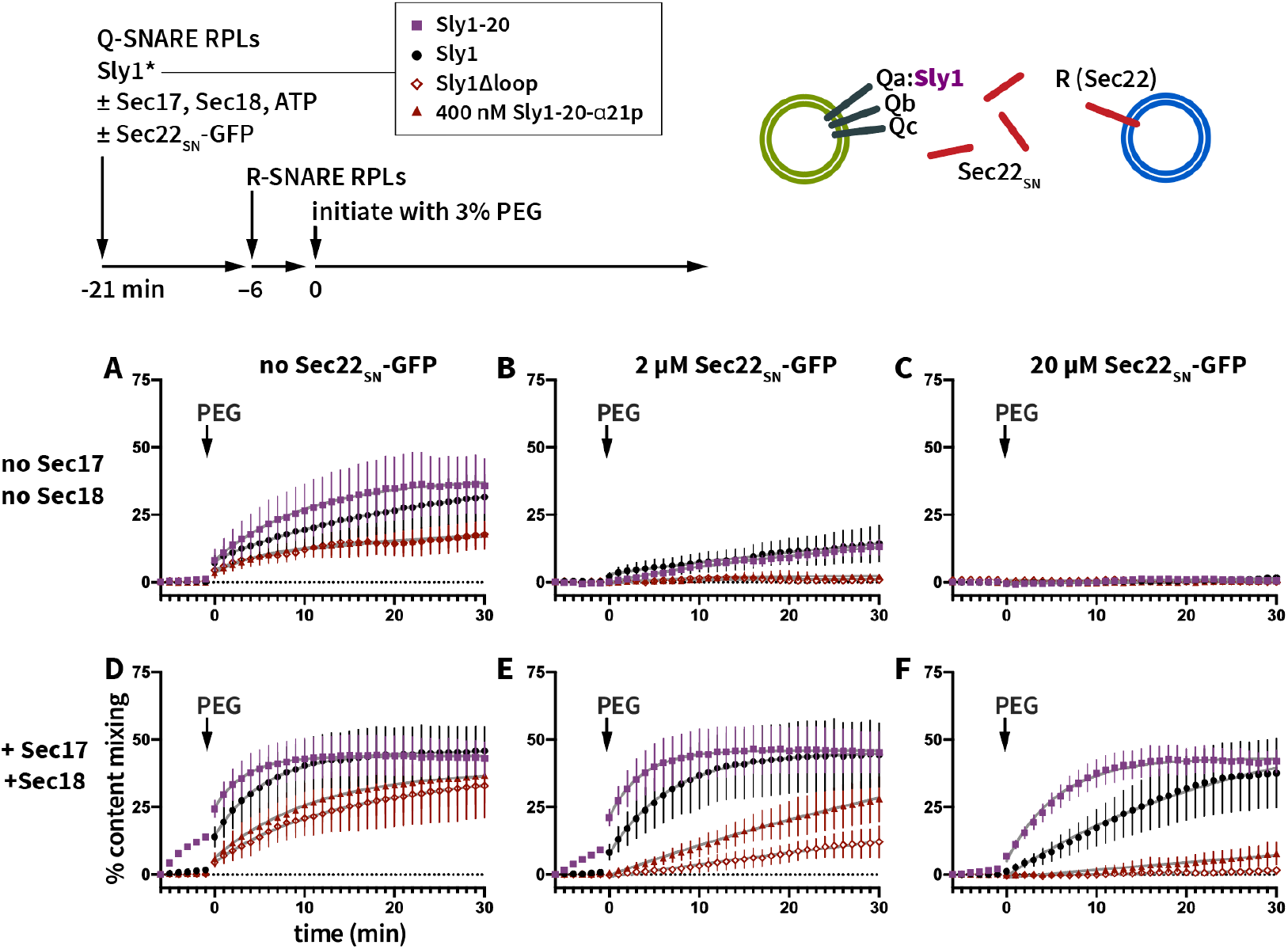
Helix α21 promotes selective formation of fusion-active *trans*-SNARE complexes. This experiment is similar to the one shown in Fig. 6, except that PEG was added to 3% final rather than 4%. At t = −21 min., Q-SNARE RPLs bearing Sed5-WT were mixed with 100 nM SLY1 variants as indicated in the legend, and either without (**A-C**) or with (**D-F**) Sec17, Sec18 (100 nM each) and Mg·ATP (1 mM), and with either 0 μM (**A,D**), 2 μM (**B,E**) or 20 μM (**C,F**) soluble Sec22_SN_-GFP. R-SNARE RPLs were added at t = −6 min. At t = 0, the reactions were initiated by addition of PEG to 3%. Points show mean ±sem of at least three independent experiments. Gray lines show least-squares fits of a second-order kinetic function.

**Supplementary Fig. S4.**
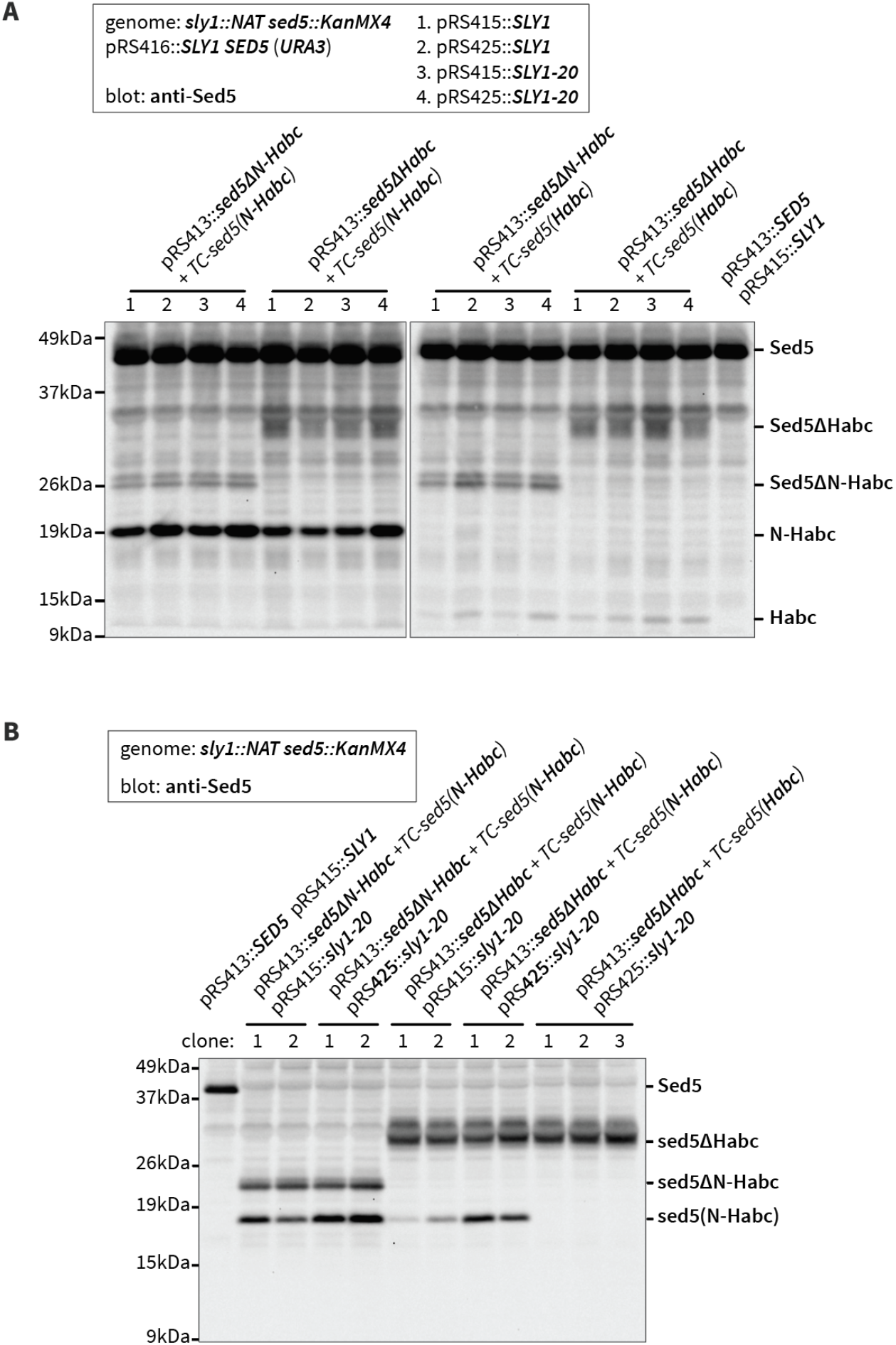
Co-expression of mutant Sed5 proteins and N-terminal Sed5 fragments. Whole-cell lysates from the indicated strains were prepared, separated by SDS-PAGE, and immunoblotted with anti-Sed5. Panel **A** shows expression in strains containing a counterselectable *SLY1 SED5* balancer plasmid. Panel **B** shows expression in strains harboring *SLY1-20* on single-copy (pRS415) or multicopy (pRS425) plasmids following ejection of the *SLY1 SED5* balancer plasmid.

